# Comparing continuum and direct fiber models of soft tissues. An ocular biomechanics example reveals that continuum models may artificially disrupt the strains at both the tissue and fiber levels

**DOI:** 10.1101/2024.09.05.610277

**Authors:** Xuehuan He, Mohammad R. Islam, Fengting Ji, Bingrui Wang, Ian A. Sigal

**Author notes:** Correspondence: Ian A. Sigal, Ph.D. Laboratory of Ocular Biomechanics Department of Ophthalmology, University of Pittsburgh Medical Center, UPMC Mercy Pavilion, 1622 Locust Street, Rm 7.382, Pittsburgh, PA, USA. 15219. www.OcularBiomechanics.org. **Disclosures:** X He: None; MR Islam: None; F Ji: None; B Wang: None; I.A. Sigal, None. Funding: Supported in part by National Institutes of Health R01-EY023966, T32-EY017271, P30-EY008098, Eye and Ear Foundation (Pittsburgh, PA), Research to Prevent Blindness (unrestricted grant to UPMC’s Department of Ophthalmology and Stein Innovation Award to IA Sigal).

## Abstract

Collagen fibers are the main load-bearing component of soft tissues but difficult to incorporate into models. Whilst simplified homogenization models suffice for some applications, a thorough mechanistic understanding requires accurate prediction of fiber behavior, including both detailed fiber-level strains and long-distance transmission. Our goal was to compare the performance of a continuum model of the optic nerve head (ONH) built using conventional techniques with a fiber model we recently introduced which explicitly incorporates the complex 3D organization and interaction of collagen fiber bundles [1]. To ensure a fair comparison, we constructed the continuum model with identical geometrical, structural, and boundary specifications as for the fiber model. We found that: 1) although both models accurately matched the intraocular pressure (IOP)-induced globally averaged displacement responses observed in experiments, they diverged significantly in their ability to replicate specific 3D tissue-level strain patterns. Notably, the fiber model faithfully replicated the experimentally observed depth-dependent variability of radial strain, the ring-like pattern of meridional strain, and the radial pattern of circumferential strain, whereas the continuum model failed to do so; 2) the continuum model disrupted the strain transmission along each fiber, a feature captured well by the fiber model.

These results demonstrate limitations of the conventional continuum models that rely on homogenization and affine deformation assumptions, which render them incapable of capturing some complex tissue-level and fiber-level deformations. Our results show that the strengths of explicit fiber modeling help capture intricate ONH biomechanics. They potentially also help modeling other fibrous tissues.

**Statement of Significance:** Understanding the mechanics of fibrous tissues is crucial for advancing knowledge of various diseases. This study uses the ONH as a test case to compare conventional continuum models with fiber models that explicitly account for the complex fiber structure. We found that the fiber model captured better the biomechanical behaviors at both the tissue level and the fiber level. The insights gained from this study demonstrate the significant potential of fiber models to advance our understanding of not only glaucoma pathophysiology but also other conditions involving fibrous soft tissues. This can contribute to the development of therapeutic strategies across a wide range of application

## 1. INTRODUCTION

Glaucoma, a leading cause of irreversible blindness worldwide [2, 3], is a progressive optic neuropathy characterized by optic disc excavation and the loss of retinal ganglion cell axons that transmit visual information from the eye to the brain [4, 5]. Clinical and experimental evidence indicates that the initial site of injury in glaucoma is the ONH, in the posterior pole [6]. Elevated IOP is one of the main risk factors for glaucomatous neural tissue damage, and every current treatment is based on lowering IOP. The mechanisms by which IOP translates into neural tissue damage remain unclear [7–9].

Understanding ONH sensitivity to IOP and thus individual susceptibility to glaucoma rests, in turn, in understanding how the tissues of the ONH region manage to bear biomechanical loads. Biomechanical support to the ONH region is provided by the collagenous connective tissues of the lamina cribrosa (LC) within the scleral canal, and the adjacent peripapillary sclera and the dura and pia maters [10]. The variability in individual susceptibility to IOP-related glaucomatous damage is thought to be due, at least in part, to differences in the mechanical behavior of the ONH tissues between individuals [11]. The desire to understand the sensitivity to IOP and susceptibility to glaucoma spurred the development of computational models that can capture the mechanical behavior of the tissues and the complex anatomy of the region. Early models simplified the tissues as linear, isotropic and homogeneous [12–17] or phenomenologically nonlinear [18]. Since collagen fibers are the primary load-bearing component of the ONH tissues, there has been great interest in developing computational models that can capture accurately the mechanical behavior of fibrous tissues. ONH models have thus advanced to incorporate inhomogeneous, anisotropic and nonlinear characteristics in some cases with fiber information derived from experiments [19–25].

Recent progress in imaging technology, specifically polarized light microscopy (PLM) [26–30] and its high speed variations [31, 32], have enabled much improved visualization of the three-dimensional (3D) organization of collagen fiber bundles in the ONH [27, 28, 33]. Utilizing this detailed information, our research group has developed a direct fiber modeling framework that accounts for the complex 3D organization and continuity of the collagen fiber bundles as well as the interactions between fiber bundles, first for a small region of the sclera [34] and later for a wide region of the posterior pole incorporating the ONH, as shown in Figure 1 [1].

**Figure 1.**
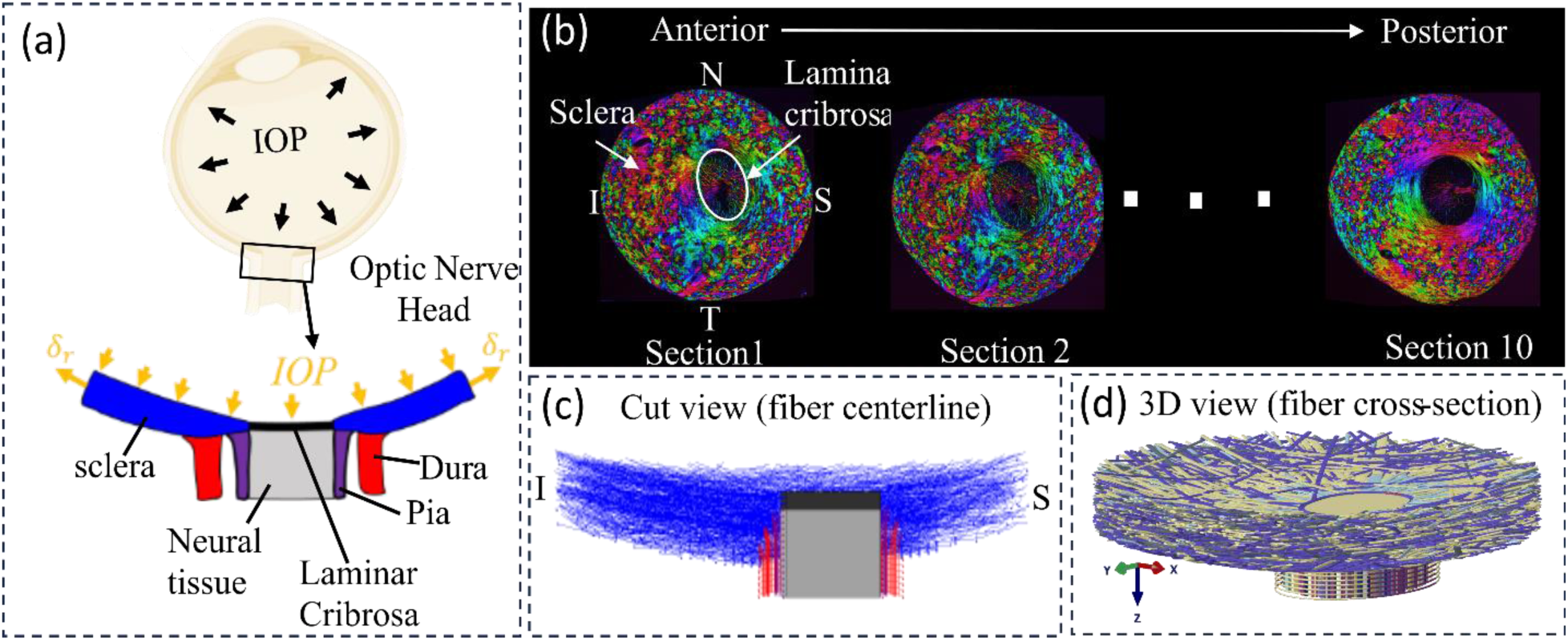
Schematic of an eye cross-section with an enlarged view of posterior sclera (blue), lamina cribrosa (black), retrolaminar neural tissue (grey), dura mater (red) and pia mater (purple) regions highlighted at the ONH. Boundary conditions included forces from IOP and displacements at the periphery. (b) Example images of serial coronal sections through the ONH of a pig eye from the anterior to posterior side. PLM was used to determine the collagen fiber orientation at each pixel [38]. Colors in the images indicate in-plane fiber orientation. Brightness indicates the strength of the signal. Low signal occurs when there is no birefringent material, no collagen, or the signal is blocked, for instance by pigment. The pig lamina cribrosa and scleral canal are elliptical with the major axis along the Nasal-Temporal (N-T) direction and the minor axis along the inferior-superior (I-S) direction. The Sections were stacked sequentially to construct a fiber model for modelling the collagenous tissues in the ONH region [1]. (c) Longitudinal cut-model view of the fiber model along the I-S direction with only bundle centerlines shown. (d) An isometric view of the complete fiber model with full bundle width displayed.

The direct fiber modeling framework employed in this model can be seen as an alternative to the conventional continuum approaches which employ constitutive models that homogenize the fibers and assume affine kinematics between individual fibers and macroscopic tissue deformation [35], without accounting for fiber-fiber interactions. Elsewhere we have shown that ignoring interweaving and fiber-fiber interactions can introduce substantial errors when estimating sclera fiber mechanical properties using inverse fitting [36]. Continuum homogenized models, and even highly simplified phenomenological models have been proven sufficient to capture gross and generic mechanical behavior of the eye. However, a thorough mechanistic understanding of the ONH region requires accurate prediction of fiber behavior, including both detailed fiber-level strains and its long-distance transmission. Our goal in this study was to compare the performance of a continuum model of ONH built using conventional techniques with the fiber model which explicitly incorporates the complex 3D organization and interaction of collagen fiber bundles [1]. To ensure a fair comparison, we constructed the continuum model with identical geometrical, structural, and boundary specifications as for the fiber model.

Specifically, we considered the fiber structure-related parameters, such as fiber dispersion, volume fraction, and orientation, as heterogeneous across the domain in the continuum model, directly calculating them based on the fiber structure reconstructed in the fiber model. We identified other material parameters following an inverse modeling approach to match the model predictions with experimentally measured average displacements at 30mm Hg. The comparison began with comparisons between the models’ predictions and experimental measures in terms of macro-level 3D tissue strain patterns and was followed by analyzing fiber-level long-distance strain transmission; corresponding results will be presented and discussed.

## 2. METHODS

### 2.1 Continuum model geometry and boundary conditions

We constructed a continuum model that aimed to mirror the fiber model introduced in [1] and shown in Figure 1, specifically in terms of geometries of the sclera, lamina cribrosa, neural tissue, pia mater and dura mater as well as the boundary conditions (Figure 2). The model was subjected to an elevated IOP of 30 mmHg. The same radial displacements (RD) used in the fiber model were applied to the continuum model’s periphery to simulate the radial tension of the sclera due to IOP. Although our goal in this work focused on the mechanical behavior of the sclera, as it was in our previous papers [1, 34, 37], the continuum model incorporated a lamina cribrosa (LC) and retrolaminar neural tissues. These provide a robust set of boundary conditions.

**Figure 2.**
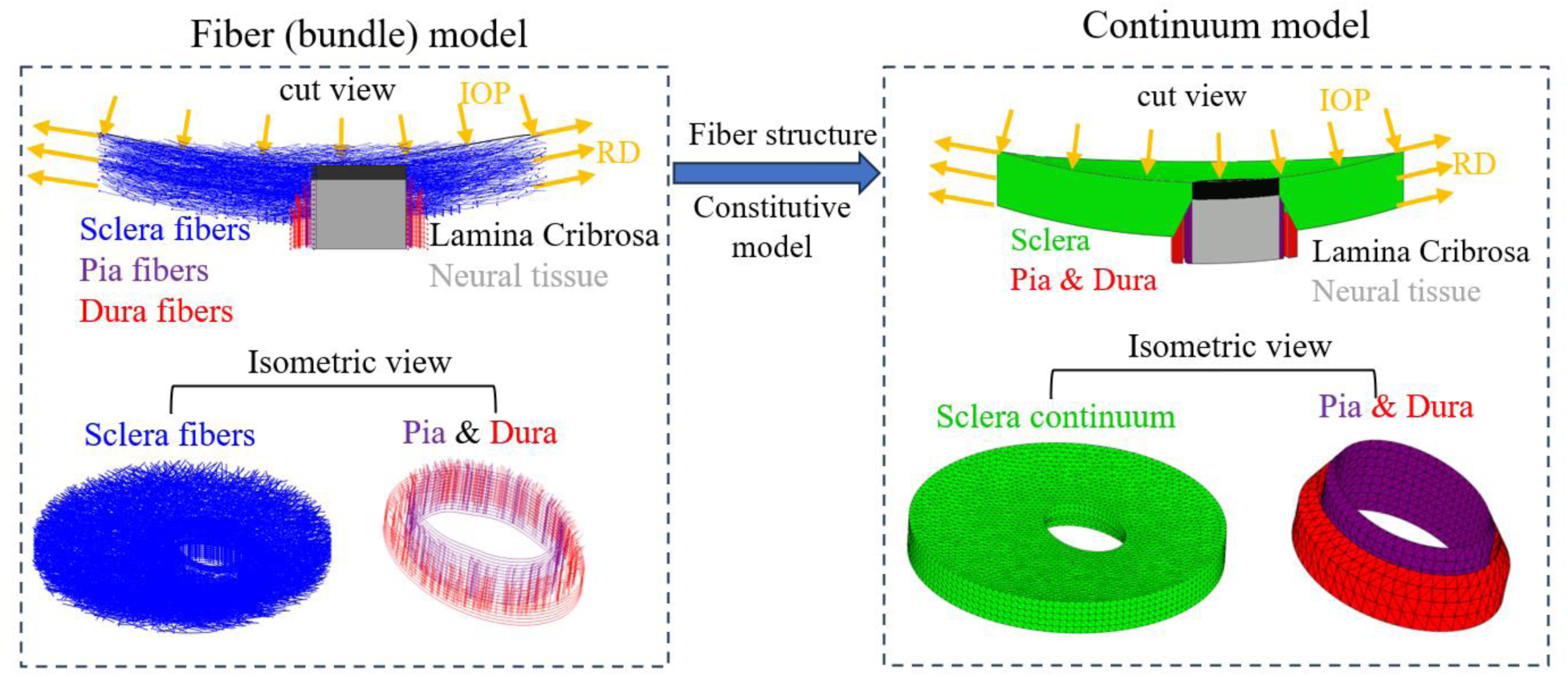
Schematic illustration of the geometry and boundary conditions of the fiber model and the continuum models. The same boundary conditions were applied to both models to simulate an inflation experiment. The continuum model was meshed using quadratic 10-nodded, tetrahedral mixed-formulation elements (C3D10H in Abaqus). A coarser mesh was selected for the peripheral sclera while a finer mesh was generated for the region where the peripapillary sclera, laminar cribrosa, neural tissue, pia mater and dura mater are located. After a mesh refinement analysis, it was decided that a mesh with ∼78,000 elements and element size ranging from 0.1mm to 0.25mm was used for the simulation.

### 2.2 Continuum model material properties

The sclera, pia mater and dura mater in the continuum model were assumed to be incompressible, anisotropic, and heterogeneous. They were characterized using the Holzapfel-Gasser-Ogden (HGO) strain energy function with one fiber family [39]. The form of strain energy function, as implemented in ABAQUS, is given by:

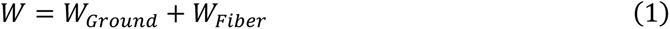

W is total strain energy density, 𝑊_𝐹𝑖𝑏𝑒𝑟_ is the strain energy density of the anisotropic collagen fibers and 𝑊_𝐺𝑟𝑜𝑢𝑛𝑑_ is the isotropic strain energy density of the non-collagenous ground matrix.

The fiber strain energy density was modeled as:

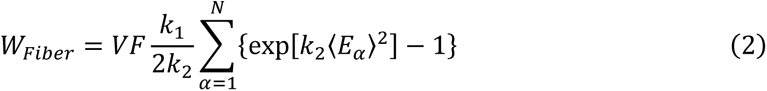

Where 𝑉𝐹 and 𝑘_1_represents the volume fraction and elastic modulus of collagen fibers. 𝑘_2_ is a material constant that governs how the stiffness of the fibers changes with the stretching of the fiber.

The strain like quantity, 𝐸_𝛼_, is expressed as:

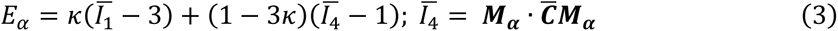

It represents the deformation of the fiber family with the mean direction, 𝑴_𝜶_, denoting as a 3D unit vector and the fiber angular dispersion, 𝜅. Here, 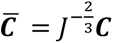 is the modified right Cauchy-Green deformation tensor; and 𝐽 is the determinant of the deformation gradient 𝑭.*Ī*_1_ is the first invariant of 𝑪̄; and *Ī*_4_ is the squared stretches in the mean fiber direction, 𝑴_𝜶_ . This model presumes that the orientation of the collagen fibers in each family is distributed rotationally symmetrically with respect to the mean preferred orientation 𝑴(𝜃, 𝜙). This rotational symmetry implies that the fiber orientation distribution is independent of the elevation angle 𝜙, i.e., 𝜌(𝑴(𝜃, 𝜙)) → 𝜌(𝜃). The parameter, κ, is then defined as follows:

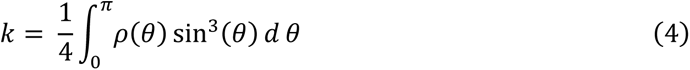

The parameter κ takes values within the range [0, 1/3]. When κ = 0, fibers are completely aligned in the mean fiber direction (no dispersion). When κ = 1/3, the fibers are randomly distributed, and the material becomes isotropic.

The strain energy density equation for the ground material was modeled as a neo-Hookean solid which has the form:

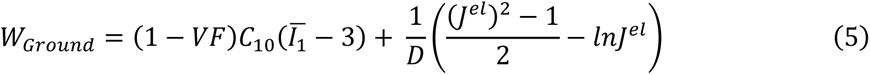

𝐶_10_is a material constant defining the stiffness of the ground substance. 𝐷 is the bulk modulus which defines the compressibility of the material. 𝐽^𝑒𝑙^is the elastic volume ratio. The incompressibility of collagenous tissue was accounted for by setting D as 0 and the utilize of mixed-formulation element type C3D10H in Abaqus.

The lamina cribrosa (LC) and neural tissue (NT) regions were modeled as linear elastic material (ELC = 0.1 MPa, ENT = 0.01 MPa), same as the fiber model.

#### 2.2.1 Identification of heterogeneous fiber dispersion, volume fraction and mean orientation

The fiber structure-related parameters of the HGO model, including fiber dispersion 𝜅, fiber volume fraction 𝑣, and mean fiber direction 𝑴, were determined on an element-by-element basis based on the fiber structure of the fiber model reconstructed from 10 PLM images of porcine ONH coronal sections as shown in Figure 1. To faithfully represent the heterogeneous fiber structural properties while maintaining the smoothness of material parameters in the tissue, in-house code was developed to calculate element-wise values of these three parameters. Briefly, for each element of the meshed sclera, pia and dura mater, neighboring elements whose center points are within 0.4mm—twice the average element size—were selected along with this element as the region of interest (ROI). Fiber segments within the ROI were then extracted. The fiber volume fraction 𝑣 was calculated as the ratio of the total fiber segments volume, considering the fiber bundle cross-section, to the total volume of the ROI. The directions of these fiber segments were fitted to a 3D π-periodic von Mises distribution to get the mean fiber direction vector 𝑴 and the concentration parameter b. The fiber dispersion 𝜅 was then related to b as:

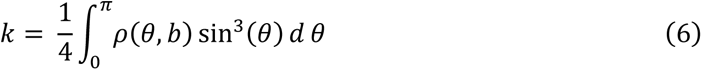

Here, 𝜌(𝜃, 𝑏) is the 2D π-periodic von Mises distribution. The derived distributions of calculated fiber dispersion 𝜅, fiber volume fraction 𝑣 and mean fiber orientation 𝑴 are shown in Figure 3.

**Figure 3.**
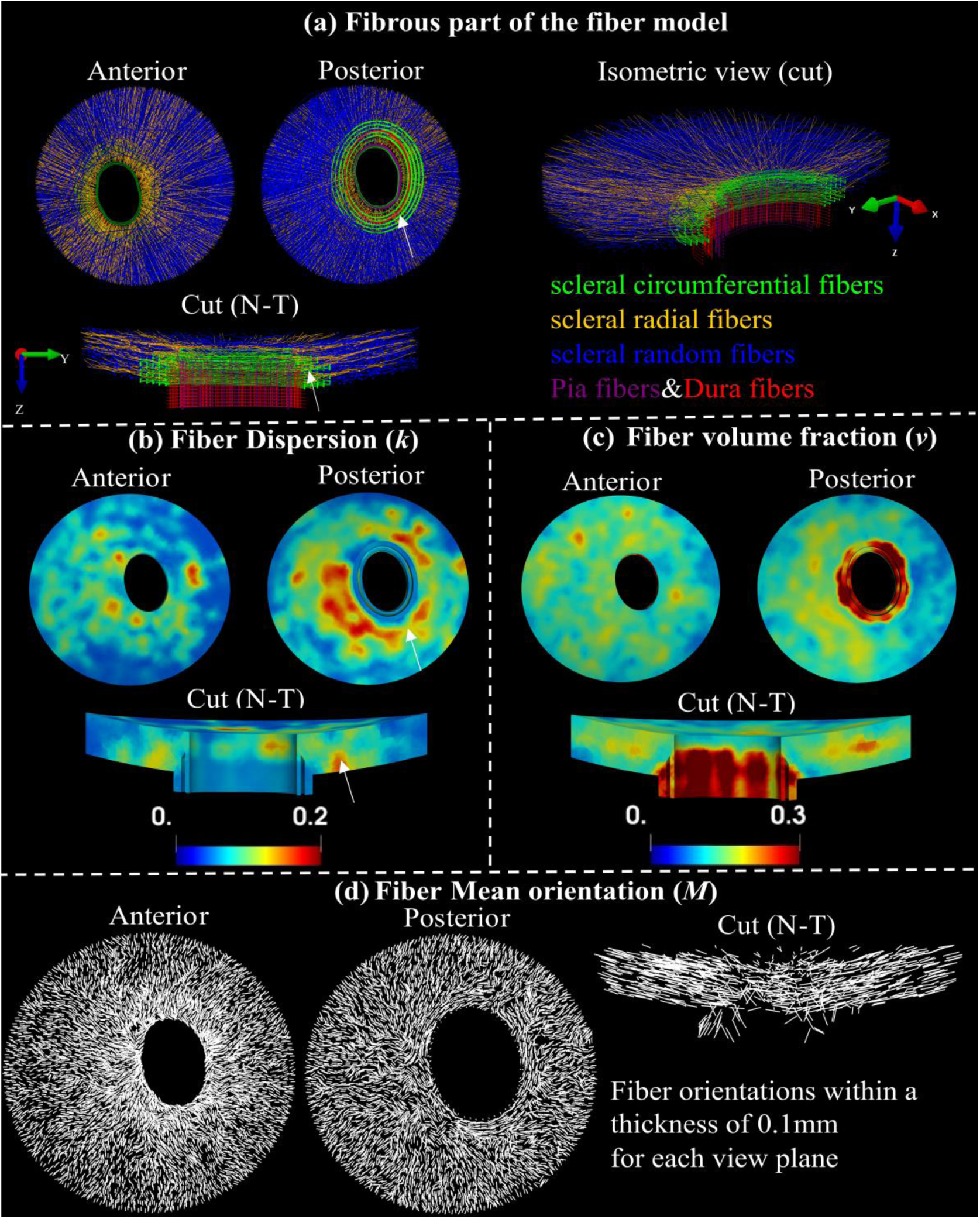
Fiber structure and contours of structure parameters. Visualization of entire fibrous regions of the fiber model with scleral random (blue), scleral radial (orange), and scleral circumferential (green), pia (purple) and dura (red) fibers in anterior view, posterior view, a sectional and an isometric cut view along the nasal-temporal (N-T) direction (a). Maps of fiber dispersion (b), fiber volume fraction (c), and mean fiber orientation (c) for these fibrous regions in anterior, posterior, and the sectional cut views. Note that the fiber dispersion here is inversely related to its degree of anisotropy. The peripapillary scleral region (indicated by the white arrows) consists of green circumferential fibers and other types of fibers (a), indicating a highly dispersed or isotropic distribution of fibers in these regions, which is associated with higher fiber dispersion (b).

#### 2.2.2 Identification of optimized matrix modulus, fiber modulus and exponential parameter

The other three HGO constitutive parameters, including the modulus of matrix 𝐶_10_, modulus of fibers 𝑢, and the exponential parameter 𝑘_2_for fibers, were regarded as homogeneous across the tissue. These parameters were identified by inversely matching the IOP-induced posterior average displacement of the nerve region and peripapillary sclera (PPS), as well as average scleral canal expansion, with those derived from ex-vivo inflation experiments documented in the literature [40]. To determine optimized parameter values, a grid table formed by 𝐶_10_ = [0.001: 0.002: 0.2], 𝑢 = [20: 10: 200], 𝑘_2_ = [50: 50: 1000] was tested. The group of values that yielded the best match of simulated responses with the experimental data were regarded as the optimized parameter values. The finite element simulation was performed using the FE solver Abaqus/Standard. Customized code and the GIBBON toolbox [41] for MATLAB v2023 [42] were used for model pre/post-processing and inverse identification of optimized parameters.

The simulation results of the continuum model with 𝐶_10_ = 0.013, 𝑢 = 120 and 𝑘_2_ = 500 produced the best match with the experimental data in terms of posterior displacements of the nerve region and PPS as well as horizontal sclera expansion. Figure 4 shows the predicted average displacements (dashed lines) with identified optimized material parameters of a porcine ONH from the fiber model (blue) and the continuum model (green) compared to the experimental measurements of different porcine eyes (circle symbols) obtained from ref. [40]. Note that the experimental displacements were measured over the ONH regions in the cross-section along the nasal-temporal (N-T) meridian direction. The same regions of the models were utilized to measure average model displacement. The shaded regions represent the standard deviation of experimental measurements from multiple eyes (n = 12). Albeit there is a slight difference of the response curves, the adjusted model predictions (solid lines) show excellent agreement with the experimental responses for all measurements. As expected, the actual model, without adjustment (dashed grey lines), does not match the experiments as well because of the difference in reference state. Please see the study introducing the fiber model for more on this topic [1].

**Figure 4.**
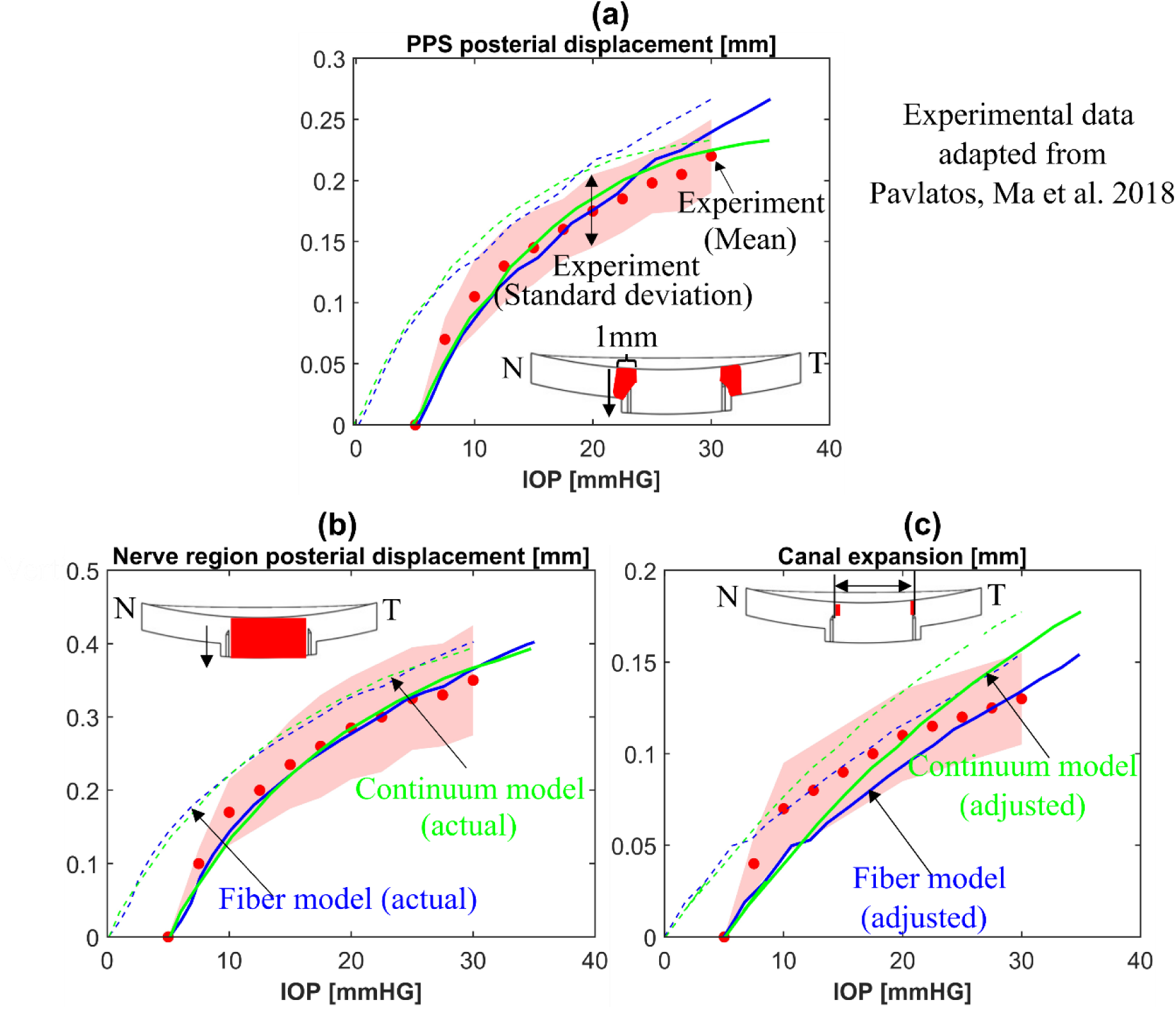
Mechanical validation of the fiber (blue lines) and continuum(green lines) models. The comparisons of (a) the mean posterior displacement of the porcine peripapillary sclera (PPS), (b) the mean posterior displacement of the optic nerve region (lamina cribrosa and neural tissue parts), and (c) the mean scleral canal expansion between the inflation experiments (circle symbols) from ref. [40] and the models (lines). Note that the specimen configurations at 5 mmHg were employed as the undeformed configuration for measuring ONH deformations in the experiments while the configuration at 0 mmHg was used as the undeformed configuration in the model. Therefore, the model responses are adjusted by 5 mm Hg in the IOP axis to account for differences in reference configurations. The shaded regions in (a)-(c) represent the standard deviation of experimental measurements for multiple porcine eyes (n = 12). The dashed and solid lines in (a)-(c) represent the actual and adjusted responses of the models, respectively. The insets in (a)-(c) indicate the regions used for calculating the mean posterior displacements and the canal expansion.

### 2.3 Comparison of strains between the continuum and the fiber models

As illustrated in Figure 5, we calculated and then compared both models in terms of the three-dimensional tissue-level strain patterns and fiber strains.

**Figure 5.**
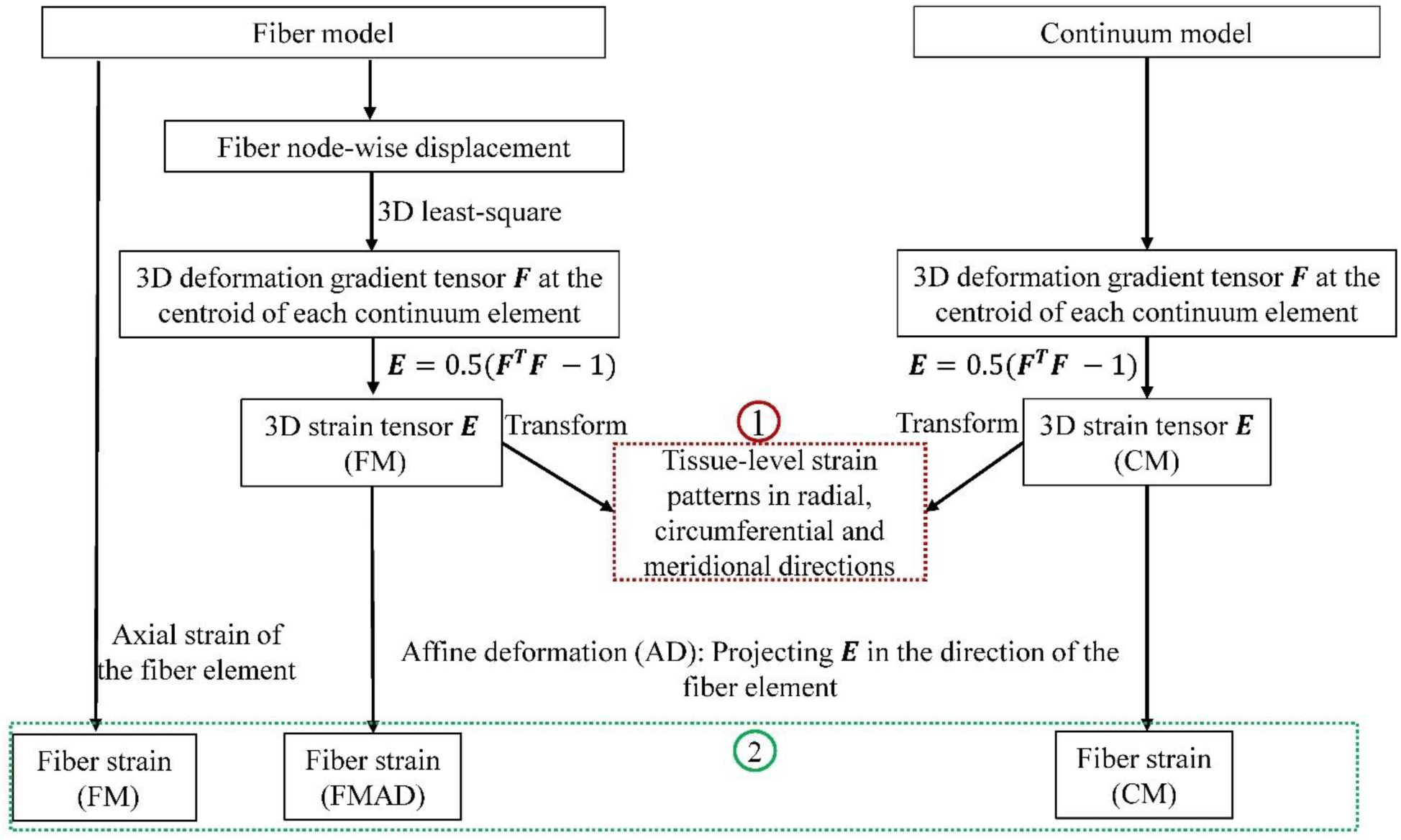
A flowchart showing the procedures in deriving 3D tissue-level strains (1) and fiber strains (2) for the fiber (FM) and continuum (CM) models. The deformation gradient tensor 𝑭 in the CM was acquired directly from the simulation. For the FM, 𝑭 was also calculated at the centroids of continuum elements to ensure consistent comparisons. The details of this calculation can be found in section 2.3.1. Tissue-level strains in radial, circumferential and meridional directions were computed by transforming the strain tensor 𝐸*E* from Cartesian coordinates to spherical coordinates, with further details on this transformation and direction conventions available in section 2.3.1. In the FM, fiber strain refers to the axial strain of the fiber element, derived directly from simulation results. For the CM, fiber strain was determined by projecting the 3D strain tensor ***E*** along the fiber element direction, following the affine deformation (AD) assumption commonly used in continuum kinematics [35]. This approach relies on the affine deformation (AD) assumption in the continuum kinematics. Employing the same AD assumption, fiber strain was also calculated based on the strain tensor derived from FM, and we refer this as the fiber strain for the fiber model with affine deformation (FMAD).

#### 2.3.1 Comparisons of three-dimensional tissue strain patterns in sclera

Several studies have reported the tissue-level 3D strain patterns of the ONH (lamina cribrosa and neural tissue parts), and peripapillary sclera (PPS) derived from experimental inflation tests [40, 43–46]. We calculated the 3D tissue strains in the radial, circumferential, and meridional directions from the continuum and fiber models at 30 mm Hg, following the procedures illustrated in Figure 5, and then compared their patterns with those reported in experimental studies.

For both models, the green strain tensor 𝑬 at the centroid of each continuum element was calculated as 𝑬 = 0.5(𝑭^𝑻^𝑭 − 1) where 𝑭 is the deformation gradient tensor. For the continuum model, 𝑭 was acquired directly from the simulation. For the fiber model, it was postprocessed based on the displacements of fiber nodes derived from the simulation. For each element’s centroid in the continuum model, neighboring fiber nodes within 400um were selected and corresponding displacements vectors in the Cartesian coordinate were denoted as 𝑈_𝑖_ (𝑖 = 𝑥, 𝑦, 𝑧) . The deformation gradient tensor 𝑭 in the Cartesian coordinate was calculated as follows:

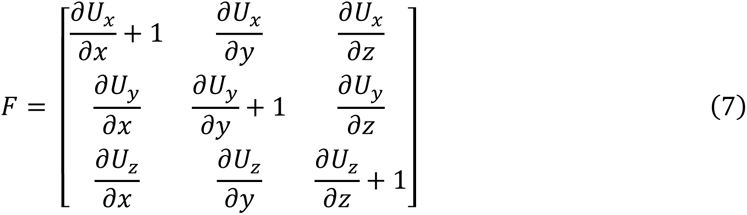

where the displacement gradients 𝜕𝑈_𝑖_⁄𝜕𝑈_𝑗_ were obtained by a 3D least-squares method [47, 48].

The derived Cartesian strain tensor 𝑬 was transformed into spherical strain tensor 𝑬_𝒔𝒑𝒉_ in the spherical coordinate via a transformation matrix T as 𝑬_𝒔𝒑𝒉_ = 𝑇𝑬𝑇^𝑇^ [45, 49]. The T is given by:

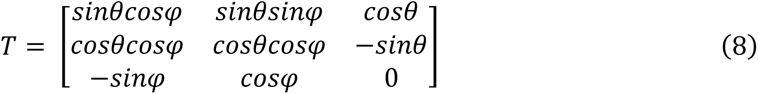

Where 𝜃 is the azimuth angle and φ is the elevational angle as shown in Figure 6. The diagonal components of derived spherical strain tensor 𝑬_𝒔𝒑𝒉_ represent the normal strain in radial 𝐸_𝑟_, meridional 𝐸_𝜑_ and circumferential 𝐸_𝜃_ direction as denoted in Figure 6, respectively.

**Figure 6.**
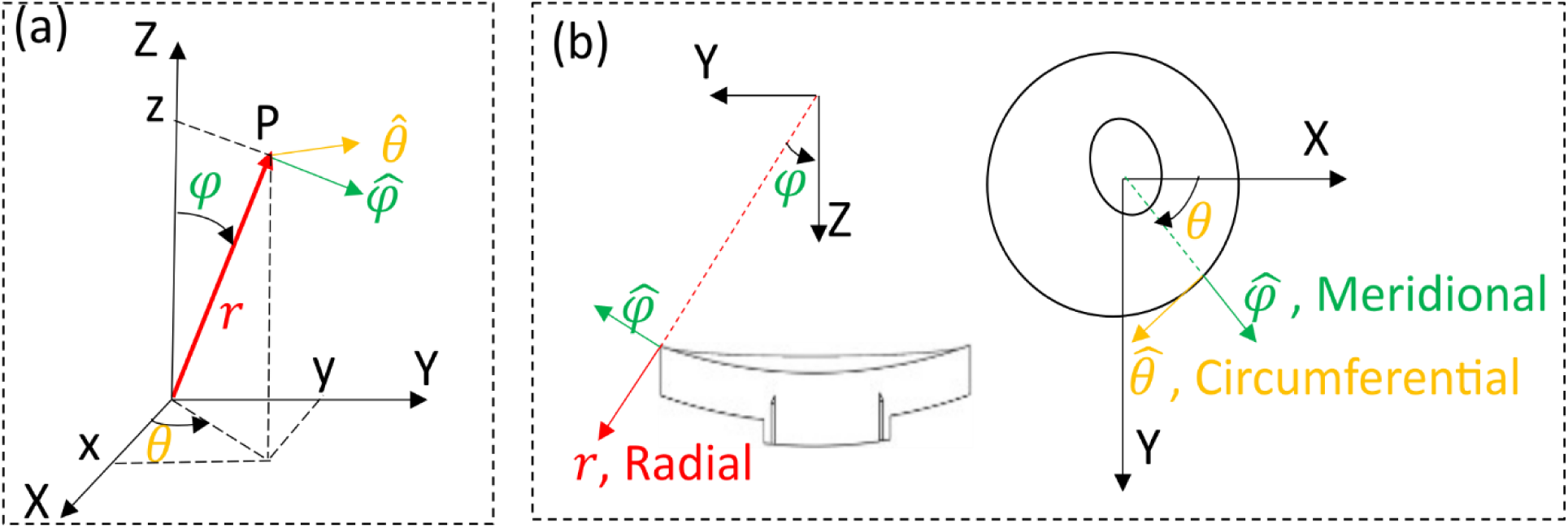
(a): Schematic of the 3D cartesian coordinate (X, Y, Z) and spherical coordinate (𝒓, 𝜽, 𝝋) systems. (b): Illustration of the radial, meridional and circumferential directions for the ONH model in cut view (N-T direction) and anterior view. The radial direction was defined to be aligned with the through-thickness direction of the ONH, which agrees with the experimental data

**Figure 7.**
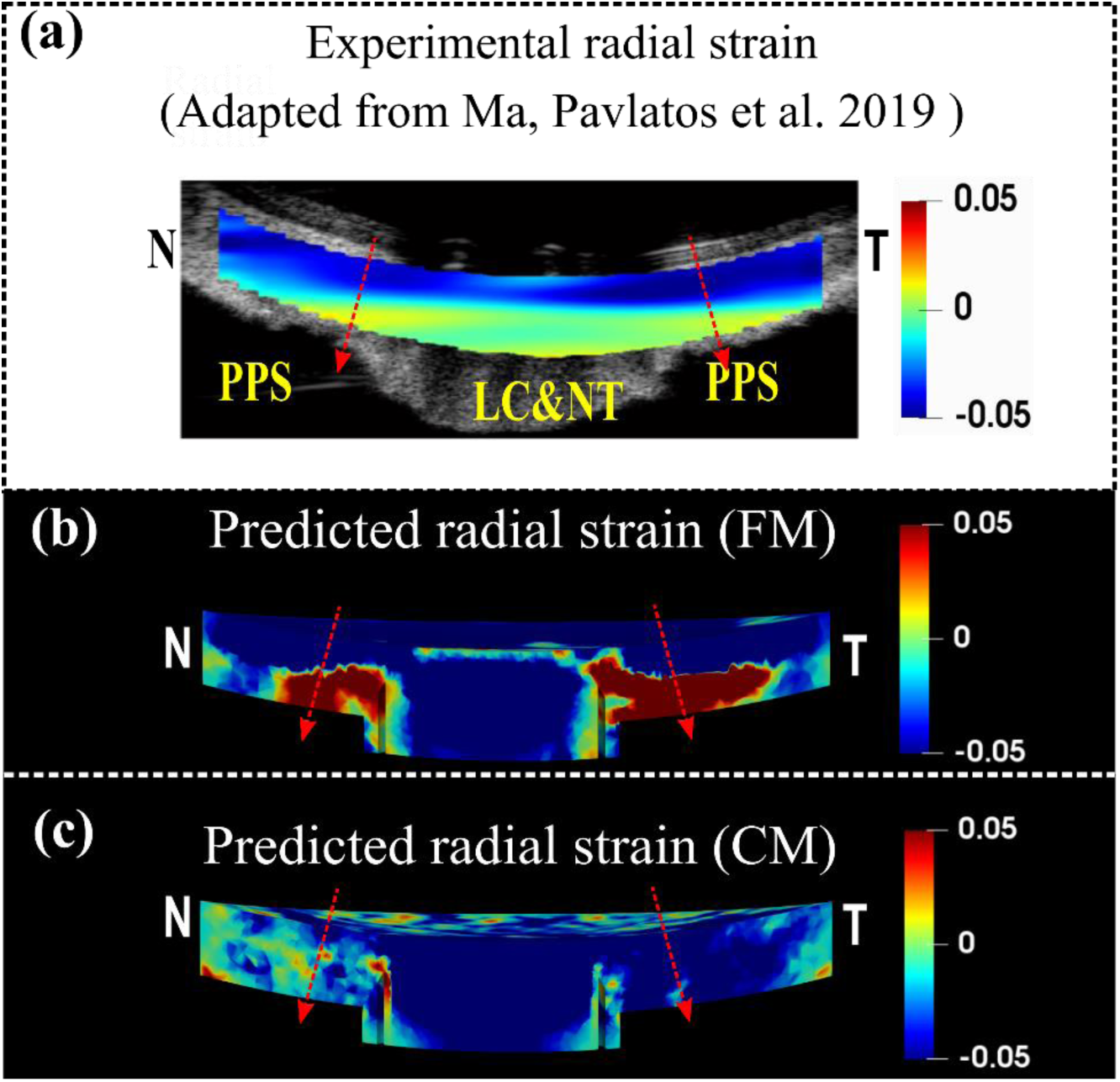
A comparison of the experimental (a) [46], the FM predicted (b) and the CM predicted (c) radial strains for the sagittal section of the optic nerve head along the Nasal-Temporal (NT) direction at 30 mm Hg. The red arrows indicate the direction of radial strain at through-thickness points, represented by corresponding dashed lines, which are oriented from anterior to posterior. The detailed direction convention can also be found in Figure 6. The experimental radial strain derived from a human eye exhibits depth-dependent variation, compressing more at the anterior and less, or stretching, at the posterior (a). Note that the fiber model faithfully replicated this pattern, but the continuum model failed to reflect it.

### 2.4 Comparison of fiber strains between the continuum and fiber models

We followed the procedures depicted in Figure 5 to calculate and compare fiber strains between the CM, FM, and FMAD at 30 mm Hg. This comparison focused on the distribution of fiber strains for fiber elements along each entire fiber in scleral fiber bundles. To quantify the degree of variation in fiber strains, we calculated the standard deviations of fiber strains along each entire fiber derived from the three models for all fibers.

## 3. RESULTS

### 3.1 Comparison of three-dimensional tissue strains between the continuum and fiber models

Predicted tissue-level radial strain patterns from the FM and CM were compared to experimentally measured patterns from a human eye [46] (Figure 5). The experimentally measured radial strains exhibited depth-dependent variability from the anterior to the posterior side, with significant compression in the anterior, whereas the posterior side of the LC, NT and PPS were less compressed or even stretched. In addition to the study shown in Figure 5. a, this pattern has also been reported in another study for human sclera tissue [45]. The FM accurately replicated the depth-dependent variation of the radial strain for the PPS, whereas the CM did not. It is important to acknowledge that the LC and NT part, modeled as a continuum in both models, did not reproduce the experimental radial strain patterns.

The FM also replicated some interesting features of circumferential and meridional strain patterns observed experimentally from the posterior side of a human sclera [44], whereases the CM failed to (Figure 8). Here, the circumferential strain seems to display a radial pattern, with bands emanating from the scleral canal and extending outward in all directions. In contrast, the meridional strain tends to follow a ring-like pattern, with large strains surrounding the scleral canal. The FM effectively captured these contrasting patterns and the pronounced near-canal meridional strains.

**Figure 8.**
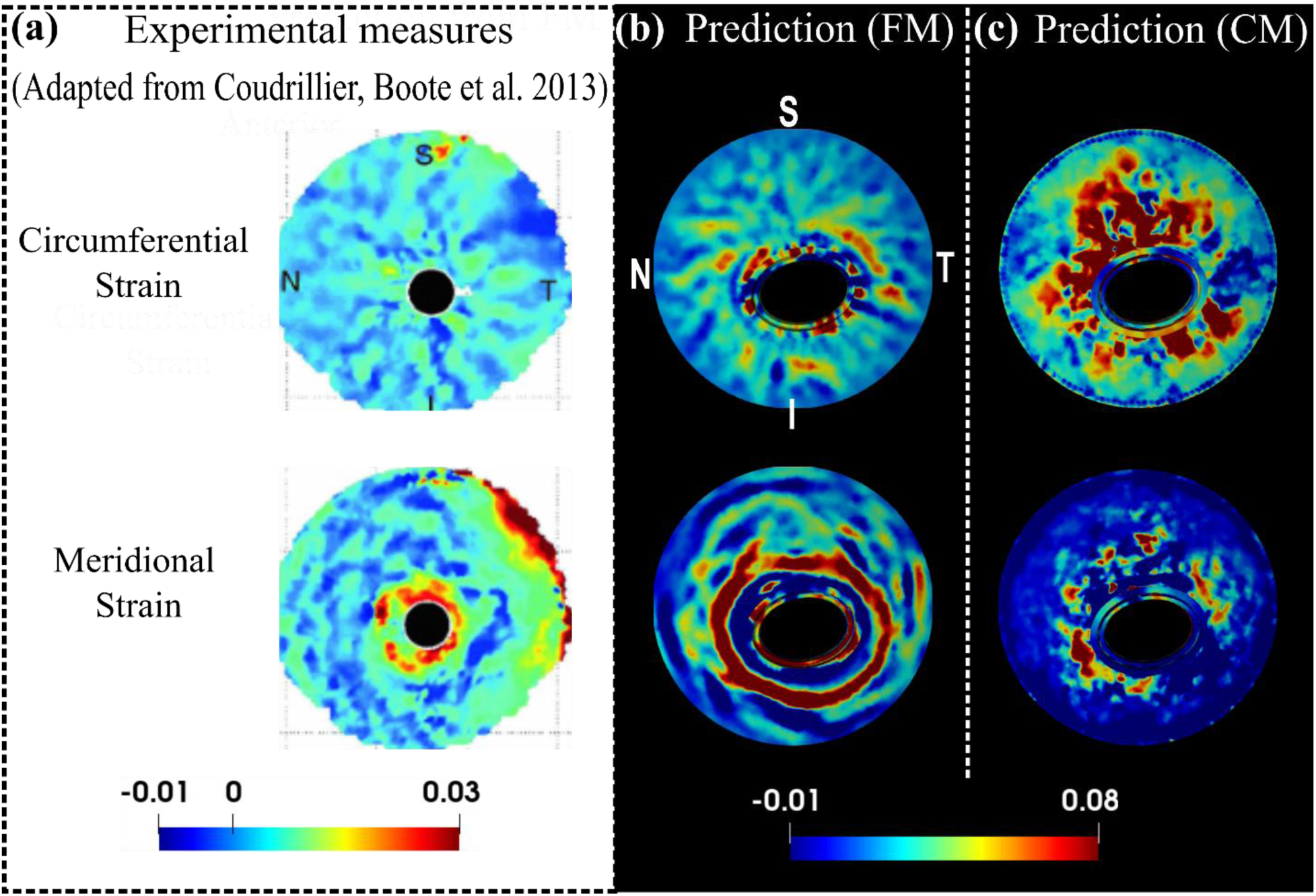
A comparison of the experimental (a) [44], the FM predicted (b) and the CM predicted (c) circumferential and meridional strain patterns from the posterior view of the ONH at 30 mm Hg. Detailed direction conventions for the circumferential and meridional strains are described in Figure 6. The plots of model predictions (b and c) were adjusted to align with the experimental plots in terms of the positions of the four quadrants: nasal (N), temporal (T), superior (S), and inferior (I). The experimental patterns, derived from a human sclera specimen [44], interestingly show contrasting behaviors: the circumferential strain radiates from the sclera canal, forming a pattern with outward-extending bands, whereas the meridional strain appears as a series of open, ring-shaped bands with high values concentrated near the canal. These contrasting patterns and pronounced near-canal meridional strains were accurately captured by the fiber model but not by the continuum model.

### 3.2 Comparison of fiber strains between the continuum and fiber models

The fiber strains derived from the FM, CM and FMAD for fiber elements along each entire fiber are shown in Figure 9. Visually, the fiber strains in the FM appear quite smooth along each fiber but vary significantly in the CM and FMAD. This observation is further supported by the results of the standard deviations of fiber element strains along each fiber for all fibers, as shown in Figure 10. Overall, the fiber strains exhibit a much lower degree of variation in the FM than in the CM and FMAD along each fiber.

**Figure 9.**
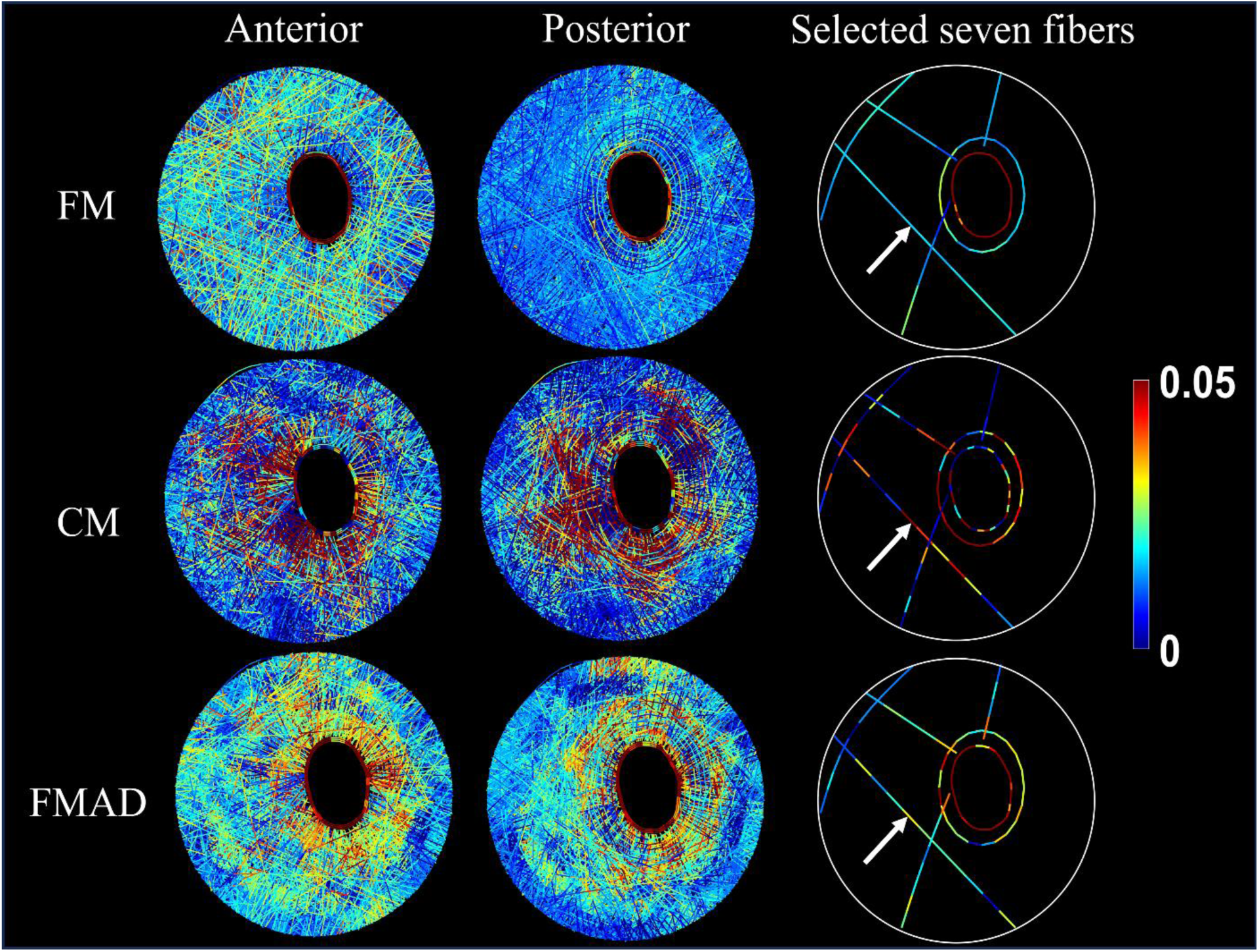
Visualization of fiber strains for all sclera fibers (anterior view in the first column and posterior view in the second column) and seven selected whole fibers (third columns) in FM, CM, and FMAD. The selection of seven fibers allows for a clearer investigation of the fiber strain changes along each fiber. Further information on the fiber strain calculation methodologies for each model can be found in Figure 5 and section 2.3.1. Take the fiber indicated by the arrows as an example. Note how the smooth strains along the fiber in the FM were disrupted in the CM and FMAD due to the enforcement of affine deformation in calculating the fiber strains. Interestingly, the fiber strains within the FM exhibit depth-dependent variation, with larger strains evident in the anterior view and smaller strains in the posterior view. This variation appears to be less pronounced or absent in the CM and FMAD models.

**Figure 10.**
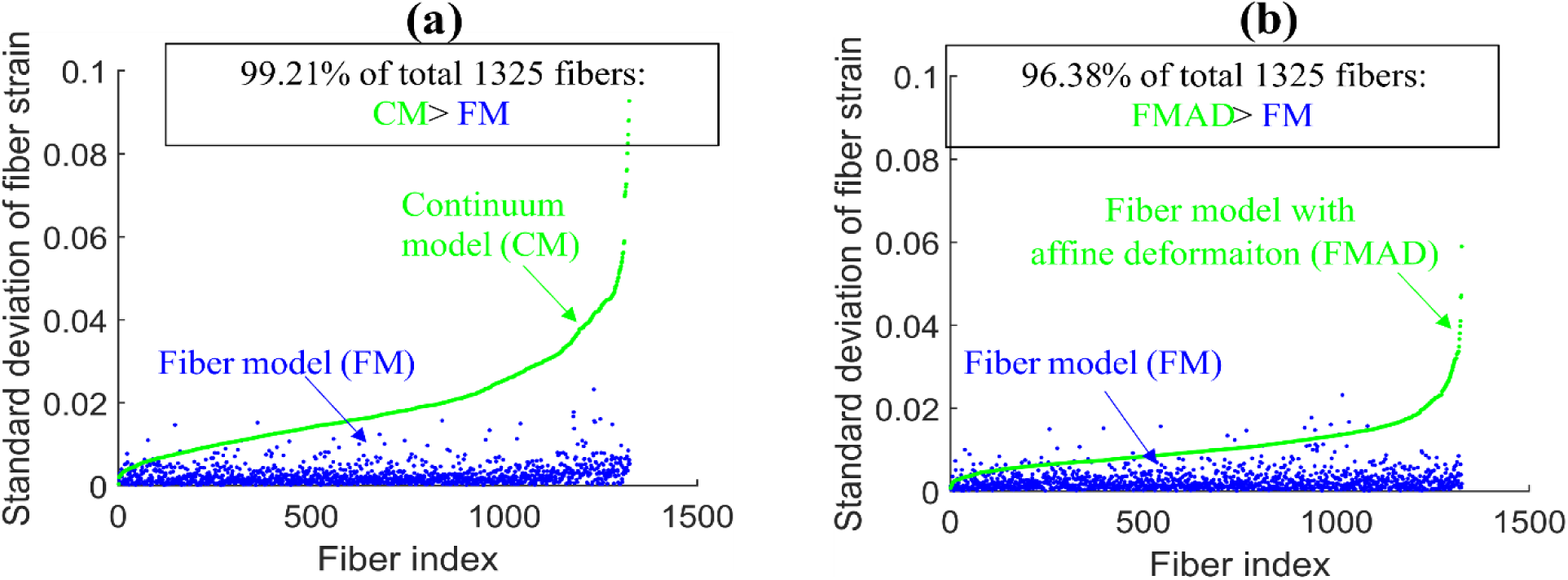
Scatterplots of the standard deviations of fiber element strains along each of the 1325 fibers derived from CM and FM in (a) and from FMAD and FM in (b). Fibers are indexed according to the standard deviation of fiber strains from the CM in (a) and FMAD in (b). The plots highlight that fiber strains from FM are notably consistent along each fiber (low standard deviations), with 97% of fibers from FM exhibiting standard deviations below 0.01.whereas those calculated enforcing affine deformation in continuum kinematics (CM and FMAD) show significantly more variability.

## 4. DISCUSSION

In this study, we employed the ONH as a test case to assess the biomechanical performance of our FM compared to a conventional CM developed using standard techniques. Our findings reveal that, while both models adequately matched the globally averaged displacement responses induced by IOP observed in experiments, they diverged dramatically in their ability to replicate specific 3D tissue-level strain patterns. Notably, the FM excelled in capturing the depth-dependent variability of radial strain, the ring-like pattern of meridional strain, and the radial pattern of circumferential strain —patterns which the CM failed to reproduce. Moreover, the FM preserved the smooth strain transmission along each fiber, in contrast to the conventional CM, which disrupted this transmission. Below, we discuss the motivation and rationale for the study as well as the significance of each finding.

Collagen fibers are the main load-bearing component of soft tissues but difficult to incorporate into models. Conventional CMs represent the collagenous fibers in soft tissues as homogenized continuum structures, which are effective for predicting macro-scale responses but lack the capacity to provide detailed fiber-level behaviors necessary to fully comprehend fibrous tissue biomechanics [22, 24, 44, 50]. To overcome these shortcomings, we have developed a FM that explicitly incorporates the complex 3D organization and interactions of collagen fiber bundles within the ONH. The motivation for this research is to highlight the significant advantages of our fiber model over conventional continuum models. By using the ONH as a test case, we aim to demonstrate that our model can more accurately predict both the intricate tissue-level strain patterns and the long-distance strain transmission along the fibers, which are crucial for advancing our understanding of fibrous tissue biomechanics and linking tissue biomechanics with cellular mechanobiology [51].

We acknowledge that the models we compare are distinct, by nature, and therefore that there will always be differences between them. Nevertheless, we contend that the comparison between them is fair, for the following reasons: Both models were developed with identical geometric and boundary specifications, ensuring a level basis for comparison When it comes to material properties, the two models account for the properties of the same fiber structure but intrinsically diverge in their approaches to representing it. The FM accurately reconstructed the fiber structure, assigning a linear stiffness to each fiber bundle. In contrast, the CM simplifies the fiber structure into a homogenized continuum and employs an anisotropic, structurally motivated constitutive model to account for the material properties. In this constitutive model, fiber structure-related parameters—such as volume fraction, dispersion, and mean orientation were directly calculated and assigned to each mesh element based on the detailed fiber structure used in the fiber model. For each element, the region of interest (ROI) is defined as twice the average element size to ensure inclusion of surrounding elements. This approach guarantees the smoothness of the material domain and replicates the local structural properties as accurately as possible. Other material parameters, including the modulus of the matrix and fibers, and the exponential parameter, were assumed to be homogeneous across the domain. These parameters were inversely identified by matching the model predictions with experimentally measured responses. It is important to note that the assumption of homogeneity for these parameters is not arbitrary but follows standard practice in these types of studies [19, 21, 23]. Furthermore, unlike previous studies that typically rely on estimated parameter values from prior research, our study enhances the accuracy and relevance of our models by employing an optimization process through inverse modeling, aligning with the method used in determining fiber stiffness for the FM [1]. However, it is crucial to recognize that despite our efforts to align the models as closely as possible, the two approaches remain inherently distinct. No modeling approach can perfectly replicate another due to fundamental differences. Thus, while our efforts ensure a fair comparison, they also highlight the unique contributions and limitations inherent in each modeling approach.

Our results demonstrated that both models performed equally well in replicating the experimentally measured nonlinear IOP-induced average displacement responses of the ONH through an inverse modeling approach, with one parameter optimized for the FM and three for the CM (Figure 4). However, significant differences emerged when it came to capturing 3D tissue-level strain patterns. The FM, despite using only one parameter to represent the linear, isotropic fiber bundles modeled as beams, was able to capture both the nonlinear IOP-induced mean displacement responses of the ONH and the intricate 3D strain patterns in the sclera. In contrast, although the CM employed six constitutive parameters and could replicate the nonlinear displacement responses, it failed to capture the 3D strain patterns of the sclera. Previous studies [40, 43] have also reported similar depth-dependent radial strain patterns in the porcine lamina cribrosa and neural tissue, showing significant anterior compression and posterior stretching. This raises the possibility that if the fibrous structures of the lamina cribrosa and neural tissue were also explicitly modeled, similar accurate results could be achieved for these regions. Further, studies have reported that the average meridional strain in the porcine ONH is greater than the circumferential strain [45, 52, 53]. This finding is consistent with the results from the FM, which showed a larger average meridional strain of 0.089 compared to an average circumferential strain of 0.0265. In contrast, the CM indicated more pronounced stretching in the circumferential direction, with average meridional and circumferential strains of 0.017 and 0.042, respectively. This can also be observed in Figure 8. However, we have not highlighted this finding as a main result of our study because it is derived from just one specimen. Nevertheless, it underscores the potential of the FM. These findings highlight that while the CM is effective for modeling large-scale tissue displacements, it is limited in its ability to represent complex 3D deformation patterns. Conversely, the FM shows considerable promise in capturing these detailed strain patterns, emphasizing its potential to provide a more accurate representation of biomechanical behavior in fibrous tissues.

In the second stage of our comparison, we focused on strain transmission along each fiber. The FM allows for direct visualization of fiber kinematics and consistently demonstrated smooth strain distribution along each fiber, as evidenced in Figure 9. In contrast, the CM exhibited substantial variation in fiber strain (Figures 9 and 10) due to its inherent kinematic assumptions. To address concerns that the observed “unsmoothness” or variation in fiber strain within the CM was due to discrepancies in tissue-level strain distributions between the continuum and FMs, we introduced the FM with affine deformation (FMAD). This model recalculated fiber strain using the same affine deformation assumption typically applied in continuum kinematics but based the calculation on the 3D strain tensor derived directly from the FM. This approach ensures that the observed strain “unsmoothness” is not merely a result of differences in the macro-level strain distributions between the models. However, even when FMAD was employed to calculate the fiber strain, the “unsmoothness” remained significant, as illustrated in Figures 9 and 10. This persistence of variation suggests that the issue extends beyond simple alignment of tissue-level strains and points to fundamental limitations of the affine deformation assumptions in continuum kinematics. This finding demonstrates the reliability of the FM and underscores the need for a thorough evaluation of the CM’s ability to link macroscopic tissue responses with cellular and sub-cellular level activities.

We recognize the substantial expertise within the field dedicated to advancing CMs of fibrous soft tissues. To bridge the gap between macroscopic observations and microscale behavior, multiscale CMs based on representative volume elements (RVEs) have been developed [54]. These models diverge from traditional constitutive material models by accounting for fiber-fiber interactions and non-affine deformations, as detailed in [55–57]. The RVE technique employs a two-scale sequential strategy, homogenizing the microscale behavior of discrete RVEs to derive the macroscale CM response. Each RVE, representing a small patch of networked fibers, influences the behavior at every integration point of the broader CM. Although innovative, the RVE method has limitations, especially due to its reliance on the continuum framework, such as the assumption of fiber independence among elements, which prevents fibers from crossing element boundaries or interacting with fibers within adjacent elements. Additionally, the homogeneous loading of RVE faces with displacements derived directly from the macroscopic solution implies an affine assumption for fiber displacements at the boundary, potentially oversimplifying the complex structures and kinematics of real tissues. Therefore, despite advancements made by the RVE method, its intrinsic limitations associated with the continuum framework persist, and we believe that the FM outperforms it, particularly in accurately capturing detailed fiber kinematics and long-range strain transmission in fibrous tissues.

It is important to acknowledge several limitations in this study. Firstly, the two models were constructed based on data from a single porcine eye specimen, while the experimental 3D strain patterns for the sclera were derived from human eyes (see Figures 7 and 8). However, consistent depth-dependent variations in radial strain patterns of the lamina cribrosa (LC) and neural tissue (NT) have been observed in both porcine and human specimens, with porcine specimens showing a much larger magnitude of anterior compression and posterior stretching [40, 43, 45, 46]. Additionally, human specimens exhibit similar depth-dependent strain patterns in the LC, NT, and PPS [40, 43]. Therefore, we hypothesize that both human and porcine sclera display similar strain patterns, with individual specimen differences more likely to affect strain magnitudes rather than the patterns themselves. This study focused on comparing the strain patterns rather than absolute values, reinforcing the relevance of the results despite the use of different species.

Secondly, as previously mentioned in the manuscript, the modulus of fiber, matrix, and the exponential parameters were assumed to be homogeneous throughout the fibrous region and were inversely identified. Assigning heterogeneous properties might improve the match of local strain patterns; however, the current settings already provide a good approximation of global displacement. If we considered these parameters as heterogeneous, defining a suitable distribution of them across the domain would be challenging. Additionally, the inverse fitting process would become extremely complicated, computationally expensive, and difficult to achieve unique solutions based on the currently used IOP-average displacement response. It remains unknown whether using experimentally measured deformation patterns as objectives for inverse modeling would succeed in accurately determining heterogeneous mechanical properties. However, even if such a match were achieved, it would mean that the CM requires much more complex inverse modeling and more material parameters than the FM to match the 3D strain patterns. Moreover, as discussed previously in the manuscript, matching macroscopic strain patterns does not resolve the ’unsmoothness’ of fiber strains under the affine deformation assumption.

Thirdly, the incompressibility assumption commonly used in modeling ONH [13, 19, 21, 22, 58–61], and other soft tissues [62–65] has been widely acknowledged. However, experiments have shown that the ONH and sclera exhibit volumetric compression under inflation [45, 66, 67], suggesting a reconsideration of this assumption. Compressibility of the tissue in the simulations could be achieved through tuning the bulk modulus D. However, this would significantly increase the computational burden. It is still unclear whether assuming compressibility would improve the match of strain patterns.

Fourthly, the lamina cribrosa and neural tissue are currently modeled as a homogenized continuum with linear and isotropic material properties in both models. Despite the sclera being modeled as anisotropic and non-homogeneous in the CM, this approach did not successfully match the experimental strain patterns. Given these outcomes, we suspect that modeling the lamina cribrosa and neural tissue as a continuum with anisotropic and heterogeneous properties could potentially improve results. Moreover, as discussed elsewhere in the manuscript, we believe that explicitly modeling the fibrous structures of these tissues, like the sclera fibers in the FM, could lead to better accuracy. This exploration will be a focus of our future work.

In conclusion, using the ONH as a test case, the FM demonstrates significant advantages over the CM in modeling fibrous tissues, particularly in its ability to accurately capture intricate 3D strain patterns and fiber kinematics, which are essential for understanding tissue biomechanics. While the FM has its limitations—such as the use of representative collagen fiber density, exclusion of the hydrated matrix, and assumptions about frictionless interactions and fiber crimp— it still represents a critical step forward in advancing biomechanical modeling. These limitations, as discussed in detail in the paper [1], provide opportunities for future refinement. Despite these challenges, we believe the insights from our study underscore FM’s potential and, more ambitiously, its necessity for further development. This model holds promise for studying the biomechanics and mechanobiology of the ONH and other fibrous soft tissue.

